# Structures of three actinobacteriophage capsids: Roles of symmetry and accessory proteins

**DOI:** 10.1101/2020.01.21.914465

**Authors:** Jennifer Podgorski, Joshua Calabrese, Lauren Alexandrescu, Deborah Jacobs-Sera, Welkin Pope, Graham Hatfull, Simon White

## Abstract

*Mycobacterium tuberculosis* and *abscessus* are major human pathogens that are part of the *Actinobacteria* phylum. Increasing multiple drug resistance in these bacteria has led to a renewed interest in using viruses that infect these bacteria for therapy. In order to understand these viruses, a course-based undergraduate research experience (CURE) program run by SEA-PHAGES at the University of Pittsburgh and HHMI has isolated, sequenced, and annotated over 3000 actinobacteriophages (viruses that infect *Actinobacteria*). Little work has been done to investigate the structural diversity of these phage, all of which are thought to use a common protein fold, the HK97-fold, in their major capsid protein. Here we describe the structure of three actinobacteriophage capsids isolated by students that infect *Mycobacterium smegmatis*. The capsid structures were resolved to approximately 6 angstroms, which allowed confirmation that each phage uses the HK97-fold to form their capsid. One phage, Rosebush, has a novel variation of the HK97-fold. Four novel accessory proteins, that form the capsid head along with the major capsid protein, were identified that show limited or no homology to known proteins. The genes that encode the proteins were identified using SDS-PAGE and mass spectrometry. Bioinformatic analysis of the accessory proteins suggest they are used in many actinobacteriophage capsids.

## Introduction

The *Actinobacteria* are a gram-positive phylum that contains many human pathogens, including *Mycobacterium tuberculosis* and *Mycobacterium abscessus*. A recent renaissance in using bacteriophages to treat multi-drug resistant bacteria has led to a handful of successful cases. The most recent being the treatment of a patient infected with multi-drug resistant *Mycobacterium absessus*[1]. The viruses (bacteriophages) that infect these bacteria contain a huge reservoir of genes with no known homologues to those found in the three domains of life. To characterize these phages and their genes, the Science Education Alliance Phage Hunters Advancing Genomics and Evolutionary Science (SEA-PHAGES) program advances a course-based research experience for early-career undergraduate students. Currently, there are 145 participating institutions, and the program collectively has isolated over 17,000 phages of Actinobacterial hosts [2,3], of which over 3000 have been sequenced and annotated.

Actinobacteriophages can be grouped together into over 120 clusters (plus a number of singletons with no close relatives) based on the nucleotide sequence of their genomes[4]. Although clustering is based primarily on nucleotide sequence comparisons and overall gene content, it is typical for bacteriophages within a cluster to have the same capsid morphology. Three morphologies are recognized in the order of *Caudovirales* that are characterized by tail morphology. *Siphoviridae* have a long non-contractile tail, while *Podoviridae* lack the tail altogether and only have the tail fibers needed for attachment to the host. The *Myoviridae* morphology is characterized by their contractile tail. Of the 2989 sequenced and annotated actinobacteriophage genomes, 92.9% are *Siphoviridae*, 6.2% are *Myoviridae* and 0.9% are of the *Podoviridae* morphology. This distribution is similar to other structurally characterized bacteriophages that infect a wide range of bacterial hosts[5,6]: whether these ratios reflect the real bacteriophage population or are an artifact from isolation methods is uncertain.

All three morphologies have the capsid head in common, which contains the double stranded DNA genome. All known structurally characterized *Caudovirales* share a conserved fold in their major capsid protein: the HK97-fold, first characterized in the HK97 bacteriophage[7]. Many bacteriophages also use accessory/ancillary capsid proteins.

Their roles are poorly understood and this is reflected in their nomenclature, which has multiple descriptive names for similar proteins. Minor capsid proteins/cement proteins are typically found embedded in the capsid head and are thought to play roles in stability. Decoration proteins are found on the outside of the capsid and experiments involving the removal of some decoration proteins show they have no effect on capsid stability or the infectivity of the phage, for example the Hoc protein of the T4 bacteriophage[8].

Despite the large number of annotated genomes, very little structural information is known about the actinobacteriophage, with almost all structural information coming from negative stain transmission electron microscopy. Of the actinobacteriophages, only one has a structure deposited in the electron microscopy database (EMDB), that of “Araucaria” that infects *Mycobacterium abscessus subsp. Bolletii*, at a resolution of 30 Angstroms (EMDB-2335)[9]. Solving structures of actinobacteriophages is an important part of understanding diversity and will lead to further insights into their evolution and provide insights into creating synthetic capsids for future bacteriophage therapy.

Here we describe the structural studies of three actinobacteriophages identified by students in the SEA-PHAGES program. Using cryo electron microscopy and mass spectrometry we have resolved the capsid heads of one *Myoviridae* and two *Siphoviridae* to approximately 6 angstroms and identified four novel accessory proteins.

## Materials and methods

### Actinobacteriophages and bacterial strains

The bacteriophages Patience, Rosebush, Myrna and their host, *Mycobacterium smegmatis* mc2155[10] were kindly provided by Graham Hatfull at the University of Pittsburgh as part of the SEA-PHAGES archive.

### Production of bacteriophages for cryo-electron microscopy

Thirty webbed plates (plates where the plaques are so numerous that they begin to touch) were made for each phage with 150 mm petri dishes using *M.smegmatis* lawns in top agar (Per liter: 4.7 g 7H9, 3.5 g BactoAgar, 0.1 v/v glycerol and 1 mM CaCl2 final) on Luria agar plates (Per liter: 15.5 g Luria broth base, 15 g Agar with the final concentrations of 50 ug/mL carbenicillin and 10 ug/mL cycloheximide). To 600 μL of stationary phase *M. smegmatis* (grown for 4 days at 37°C at 250 rpm in Middlebrook 7H9 liquid medium) enough phages were added to ensure a webbed plate (note: this was empirically determined with each batch of phage). Phage and bacteria were incubated at room temperature for 20 minutes to allow infection. 10 mL of top agar was then added and quickly poured onto agar plates. Patience and Rosebush were incubated at 37°C while Myrna was incubated at 30°C. Plates were incubated for 36-48 hours to allow optimal plaque formation. 10 mL of Phage Buffer (10 mM Tris-HCl pH 7.5, 10 mM MgSO_4_, 68 mM NaCl, 1 mM CaCl_2_) was then added to each plate and incubated for 4 hours at room temperature to create a phage lysate. The lysate was then aspirated from the plates and pooled together before being centrifuged at 5500 xg for 10 minutes at 4°C. The supernatant was moved to a fresh tube and the phage particles precipitated with the addition of NaCl (1 M final concentration) and PEG-8000 (10% w/v final concentration) and mixed overnight at 4°C. The phage were pelleted by centrifugation at 5500 xg at 4°C for 10 minutes and the supernatant discarded. The phage in the pellet were then resuspended in 10 mL of phage buffer by gentle rocking overnight at 4°C. The new phage lysate was then centrifuged at 5500 xg for 10 minutes to remove any debris. 8.5g of CsCl was added to the 10 mL of phage lysate and the density adjusted to a refractive index (h_D_) of between 1.3087 and 1.382 (the final density was 1.5 g/mL). The CsCl/phage solutions were then centrifuged at 40,000 xg for 16 hours and the phage band removed (it was roughly in the middle of the gradient) with a syringe and needle and stored at −80°C.

### Production of bacteriophages for mass-spectrometry

Rosebush was propagated in its host, *Mycobacterium smegmatis* mc2155 following previous methods and media[11]. The phages from the SEA-PHAGES archive were diluted and plated out onto *M. smegmatis* lawns. A single plaque was picked with a sterile pipette tip and serially diluted and plated onto *M. smegmatis* lawns. The concentration that gave webbed plates (plates where the plaques were so prevalent that they touch each other) was used to make 20 webbed plates for each bacteriophages. Plates were then flooded with 5 mL of phage buffer and incubated at room temperature overnight. Approximately 3 mL of phage buffer was then recovered from each plate (termed the phage lysate). Phage were pelleted by centrifugation at 82705 xg (25,000 rpm) for 4 hours using an SW28Ti rotor (Beckman Coulter). Phage pellets were gently resuspended overnight in 1 mL of phage buffer at 4°C with shaking. Phage were then purified using cesium chloride isopycnic purification. The 1 mL of phage was added to 6 mL of phage buffer and 5.25 g of cesium chloride (final density of CsCl was 1.5 g/mL). The gradient was then centrifuged in an S50ST rotor (Thermofisher scientific) at 103,836 xg (40,000 rpm) for 18 hours. A single phage band could be seen about half-way down the tube. The band was extracted using a needle and syringe by piercing the side of the tube. Typically 1 mL of phage was recovered. Phage were then dialyzed against phage buffer and pelleted again to concentrate the phage before being resuspended in 50 μL of phage buffer.

### Preparation of bacteriophages for cryo-electron microscopy

20 μL of each bacteriophages from the SEA-PHAGES archive (concentration unknown) was dialyzed into phage buffer (10 mM Tris-HCl, pH 7.5, 10 mM MgSO_4_, 68 mM NaCl and 1 mM CaCl_2_) using a Tube-O-Dialyzer Micro (G-Biosciences) with a 50 kDa MWCO. 3 μL of the dialyzed phage was used for uranyl acetate negative stain electron microscopy using an FEI Tecnai G2 200 keV TEM. 5 μL of Patience and Rosebush were mixed together in a 1.5 mL Eppendorf tube as were 5 μL of Myrna and Rosebush in a different 1.5 mL Eppendorf tube. These mixtures are referred to as the multiplexed phage.

### Preparation of cryo-electron microscopy grids

5 μL of multiplexed phages were pipetted onto a C-flat 2/1-2C (Protochips, 2 μm hole, 1 μm space) cryo-electron microscopy grid using a Vitrobot mk IV (FEI). Grids were blotted for 5 seconds with a force of 5 (a setting on the Vitrobot) before being plunged into liquid ethane.

### Cryo-electron microscopy

Data was collected on a 200 keV Talos Artica at the University of Massachusetts medical school. The following conditions were used. Magnification: 36000. Pixel size: 1.1 Å. 200 keV. Exposure time: 1.59787 seconds. Number of frames: 32. Dose per frame: 0.550074 e-/Å2. Defocus range: was approx. −0.5 to −2.5 μm in 0.5 μm steps. A total of 996 images were taken for the Patience and Rosebush grid while 1200 images were taken for the Rosebush and Myrna grid.

### Cryo-electron microscopy data analysis

Relion 3.0.4[12] was used for bacteriophage capsid reconstructions. Data were 2x binned so that the pixel size was 2.2 Å to speed up computation. Box size was 400 pixels (880 Å) with a mask diameter of 800 Å. 1000 particles were manually picked for 2D classification to create references for auto-picking. The Relion auto-picking was then used to pick all the phage particles. After extraction the particles were subjected to 2D and 3D classification with the 2D classes used to create the 3D initial model using the Relion software. The different capsids were clearly separated into different 3D classes. Each class was used separately for refinement using I1 icosahedral symmetry. Masks were created using the Relion software and used in the final Post-processing step in the Relion software. CTF refinement and Bayesian polishing were not performed. Figures for this paper were prepared using Chimera[13] and ChimeraX[14]. The Segger[15] function in Chimera was used to segment out the different proteins.

### Bioinformatics

The ITASSER[16,17], HHPRED[18], BLAST-P and PSI-BLAST[19] online servers were used with the default settings. The following databases were used in HHPRED: PD B_mmCIF7 0_28_Nov, PDB_mmCIF30_28_Nov, SCOPe70_2.07, ECOD_ECOD_F70_20190225, COF_KOG_V1.0, Pfam-A_v32.0, NCBI_Conserved_Domains(CD)_v3.16, SMART_v6.0, TIGRFAMS_v15.0 and PRK_v6.9. With BLAST-P and PSI-BLAST the Non-redundant protein sequences (nr) database was used.

### Mass spectrometry

#### In-gel digestions, tryptic peptide extraction and desalting

Concentrated phages were boiled in 2x SDS-PAGE loading buffer and run on a 10% denaturing SDS-PAGE. Protein ladder used was precision plus protein dual color standard (Bio-Rad). After Coomassie staining, bands were excised with a clean scalpel blade. These gel bands were destained using 40% ethanol, 10% acetic acid in water. The gel bands were diced, equilibrated to pH 8 in 0.1 M ammonium bicarbonate, and subject to Cys reduction and alkylation using 10 mM dithiothretol in 0.1 M ammonium bicarbonate (1 hr at 37°C) and 55 mM iodoacetamide in 0.1 M ammonium bicarbonate (45 min at 37°C in the dark), respectively. The gel bands were dehydrated using acetonitrile, dried to completion in a Speedvac concentrator and resuspended in a 12.5 ng/μL trypsin solution in 0.1 M ammonium bicarbonate for 45 min on ice. The supernatant was removed and replaced with 0.1 M ammonium bicarbonate. Proteolysis continued for 16 hr at 37°C on a thermal mixer. The following day, tryptic peptides were extracted using consecutive hydration and dehydration steps using 0.1 M ammonium bicarbonate and 50% acetonitrile in 5% formic acid. Two final hydration and dehydration cycles were conducted using 0.1 M ammonium bicarbonate and 100% acetonitrile. Pooled peptide solutions were dried in a Labconco speedvac concentrator and resuspended in 0.1% formic acid in water. Peptides were desalted using Pierce C18 desalting spin columns using manufacturer’s protocol.

#### Peptide and protein identification using mass spectrometry

Dried and desalted peptides were resuspended in 0.1% formic acid in water and analyzed using nanoflow ultra-high performance liquid chromatography (UPLC) coupled to tandem mass spectrometry (MS/MS) using a Dionex Ultimate 3000 RSLCnano UPLC system and Q Exactive HF mass spectrometer (Thermo Scientific). Peptides were separated using a 1 hr reversed-phase UPLC gradient over a 75 μm x 25 cm Easy Spray PepMap C18 analytical column and directly ionized into the Q Exactive HF using positive mode electrospray ionization. A high resolution MS1 and data-dependent Top15 MS/MS acquisition method was used. All raw data were searched against the Uniprot Mycobacterium phage Rosebush reference proteome (accessed 20191017, updated 20190725) using MaxQuant v1.6.0.1 and default parameters plus the following additional settings: N-terminal Gln to pyro-Glu variable modification and minimum 5 amino acids/peptide.1 Identifications were filtered to a peptide- and protein-level 1% FDR using a decoy search and uploaded into Scaffold 4 (Proteome Software) for visualization and further analysis.

## Results

### Phages, hosts, and sources

The three phages were isolated from a number of different countries and environments and have been documented previously. All were isolated using *Mycobacterium smegmatis* mc2155 host. See Table 1 for information about the phage. Phage **Patience**[4,20] was isolated in Durban, South Africa at the Nelson Mandela School of Medicine (NCBI accession number: JN412589). Relative to the GC content of the host genome (67.4%), Patience has a relatively low genome GC content (50.3%), which has been suggested to mean that it has moved from a different host (with a lower GC content than *M. smegmatis*) relatively recently[21,22]. Patience is one of two members of Cluster U and is lytic. Phage **Rosebush** was isolated from the Bronx Zoo, Bronx, NY, USA[22–24] (NCBI accession number: AY129334). Rosebush is part of the Cluster B2, which has twenty-seven members, and is lytic. Phage **Myrna**[4, 24–29] was isolated in Upper Saint Clair, PA, USA (NCBI accession number: EU826466). Myrna is one of two members of Subcluster C2, and is lytic.

**Table 1.** Basic information about the three actinobacteriophage in this paper.

| Name | Number of genes | Cluster | Number of phage in cluster | Genome length (bp) | Genome GC content (%) | Life cycle | Host | Morphology |
| --- | --- | --- | --- | --- | --- | --- | --- | --- |
| Myrna | 229 | C2 | 2 | 164602 | 65.4 | Lytic | *M. smegmatis* mc<sup>2</sup>155 | Myoviridae |
| Patience | 109 | U | 2 | 70506 | 50.3 | Lytic | *M. smegmatis* mc<sup>2</sup>155 | Siphoviridae |
| Rosebush | 90 | B2 | 27 | 67480 | 68.9 | Lytic | *M. smegmatis* mc<sup>2</sup>155 | Siphoviridae |

### Virion morphologies

All the viruses described here are double stranded DNA tailed bacteriophages of the viral order *Caudovirales* (Figure 1). Patience and Rosebush show the *Siphoviridae* morphology (long non-contractile tails), while Myrna has *Myoviridae* morphology (contractile tail).

**Figure 1.**
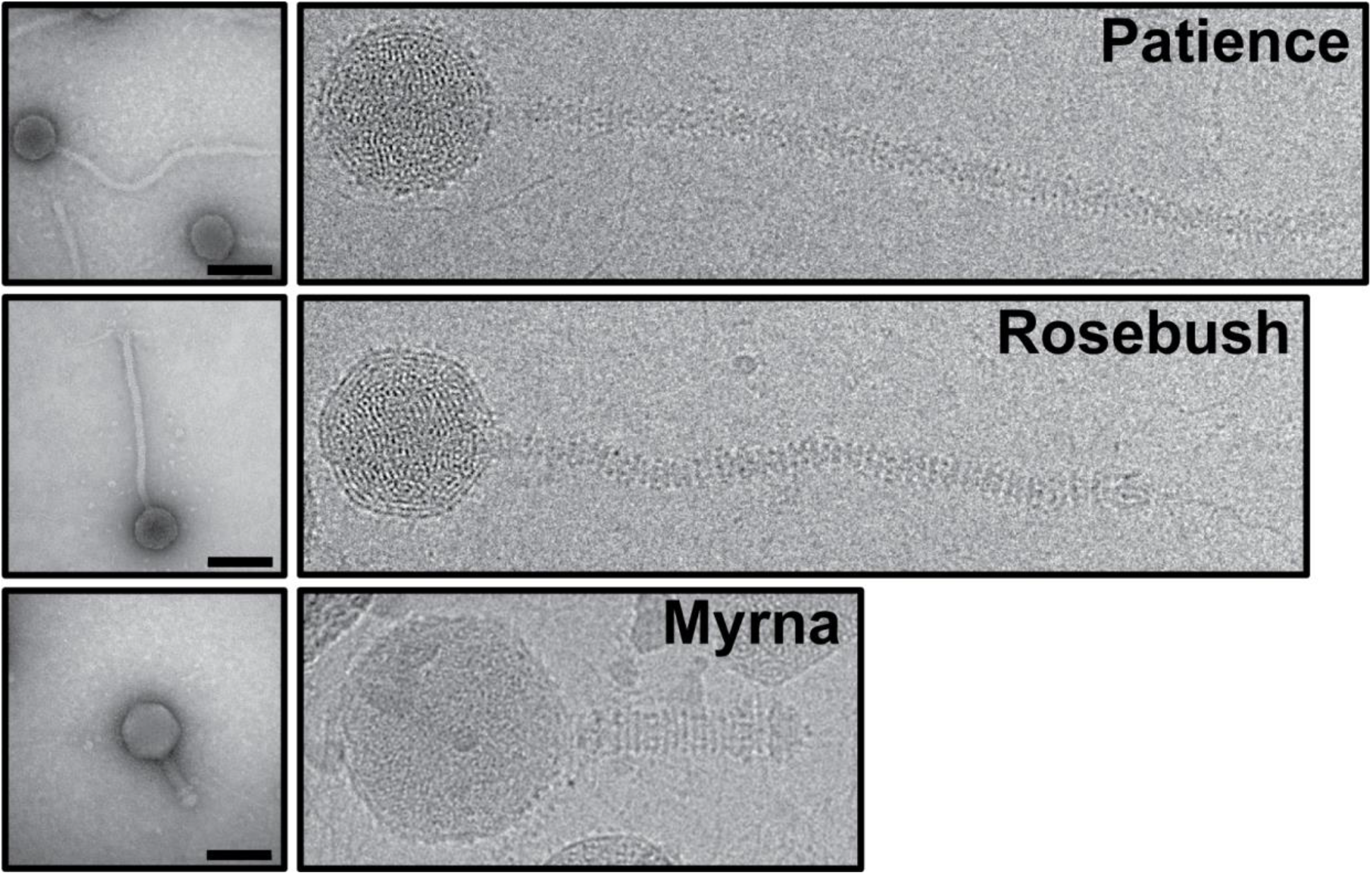
Electron microscopy of the actinobacteriophage. On the left are representative negative stain images of the three actinobacteriophages Patience, Rosebush and Myrna, while the right shows representative cryo-EM images. Scale bar in the negative stain micrographs is 100 nm. The cryo-EM images are all at the same magnification (36,000x) and their sizes are directly comparable to one another.

### Capsid morphologies and accessory proteins

The three bacteriophages were multiplexed onto two grids with Patience and Rosebush on one grid and Myrna and Rosebush on another (Figure 2 shows the typical work-flow). The rationale was that this method would allow for more structures from less microscope time. We hypothesized that the Relion software would separate the particles at either the 2D or 3D classification stage. All the particles were picked with no attempt to choose specific phage particles, for example either just Patience or just Rosebush. 2D classification had some success in separating out the different phage particles but it was not clear cut, with some ambiguous orientations of phage particles. Therefore, all “good” 2D classes, those with very clear features in the 2D class images, were selected and used in 3D initial model generation and 3D classification. It was clear in the 3D classes that the phages had been separated successfully, with the decoration proteins (described later) of Patience and Rosebush clearly recognizable. To rule out chimeric reconstructions, for example where Rosebush and Patience particles are used to make the cryo-EM map, Rosebush was added to both multiplexed grids. In both cases Relion classified the Rosebush particles separately and the final map was identical. The three bacteriophage structures were successfully resolved from the datasets (Figure 3).

**Figure 2.**
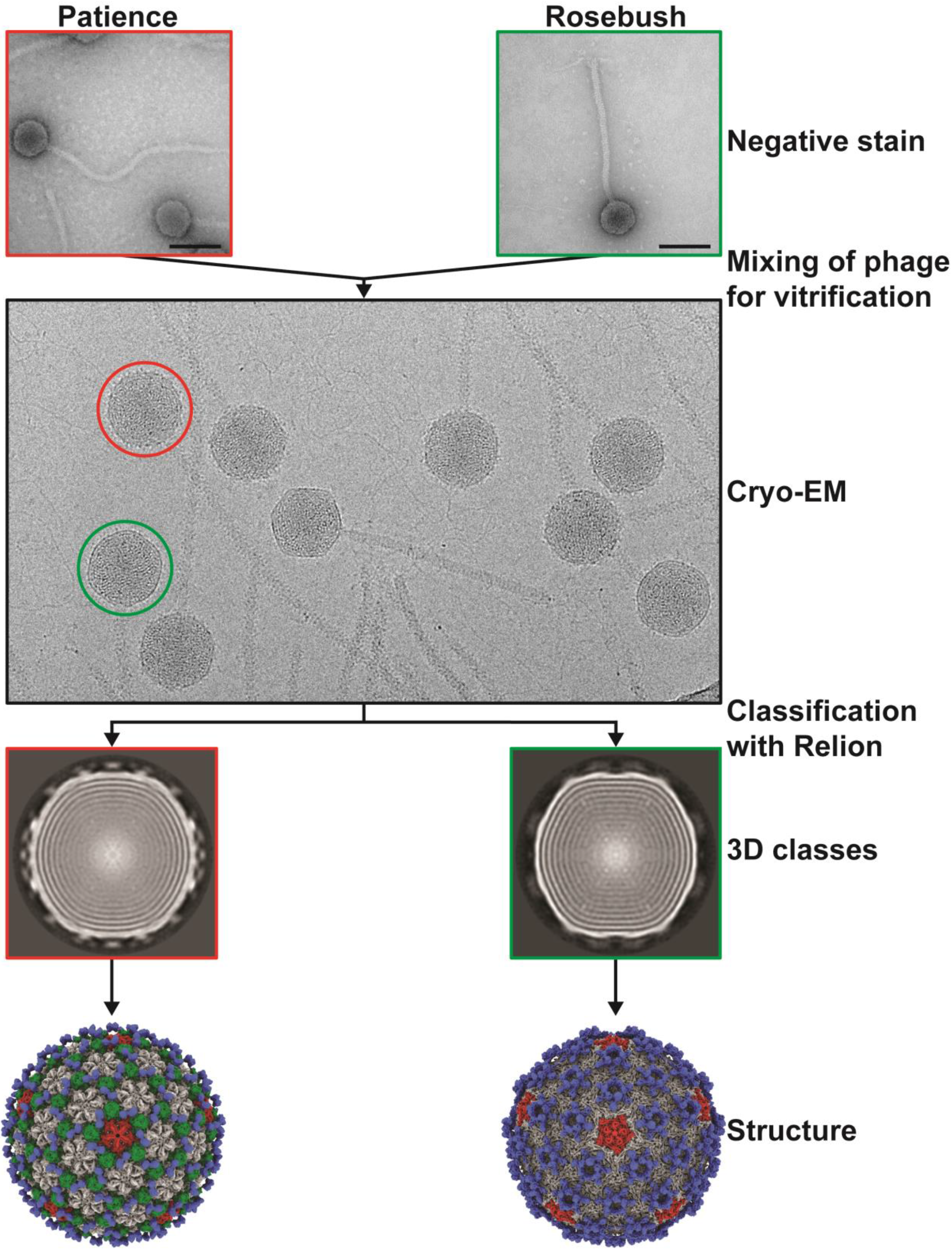
Multiplexing phage samples. Diagram showing the basic method by which the phages were multiplexed on a single cryo-EM grid and used for data analysis.

**Figure 3.**
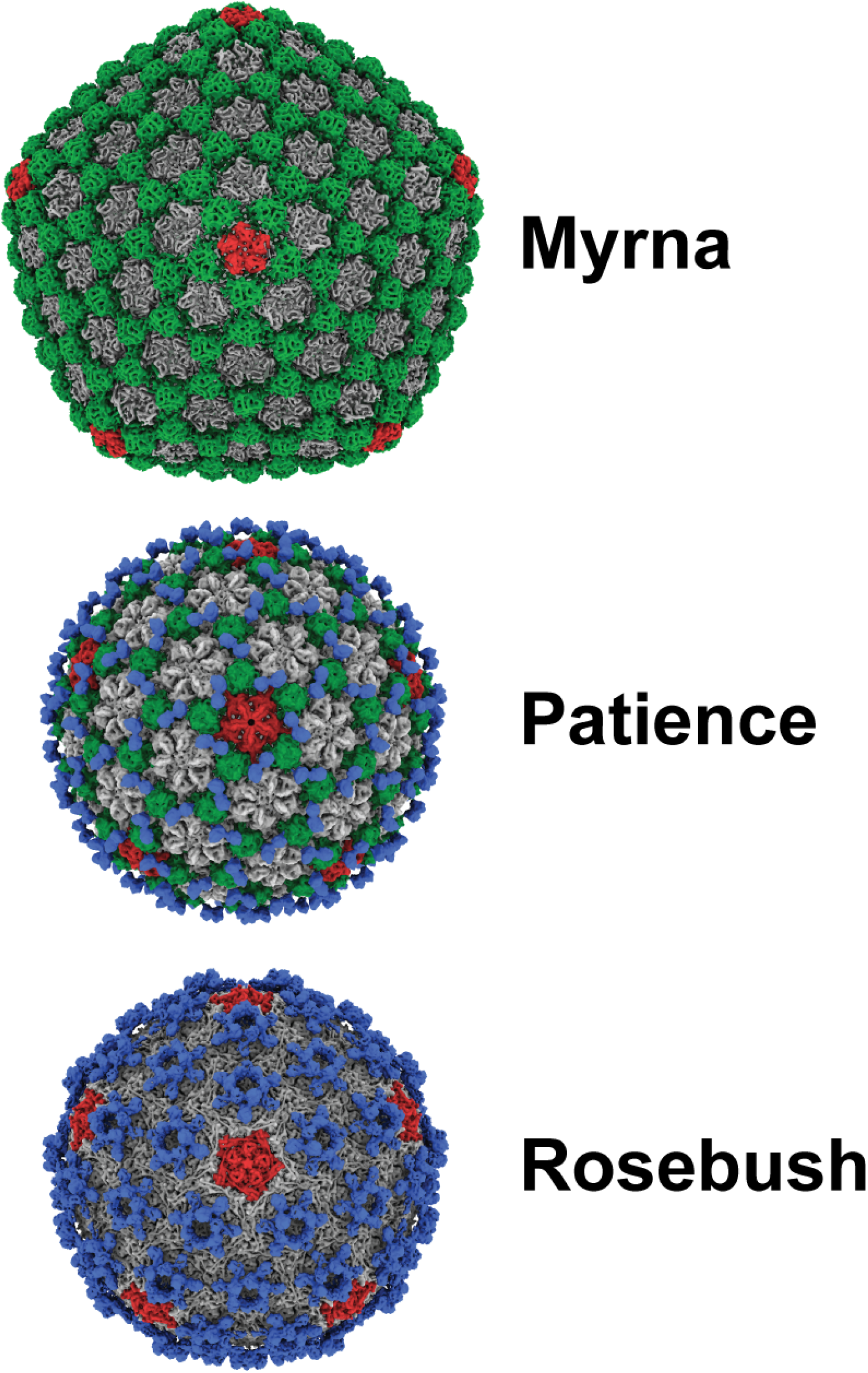
Capsid structure of the actinobacteriophages. Myrna (7.7 Å resolution from 2268 particles), Patience (5.9 Å resolution from 3461 particles) and Rosebush (6.7 Å from 3489 particles) cryo-EM maps. Pentamers are colored red, minor capsid proteins green and decoration proteins blue. Hexameric capsid proteins are grey in color. Cryo-EM maps in this figure and following figures were displayed using the ChimeraX software[14].

**Patience** shows pseudo T=7 laevo capsid organization, as is common among many dsDNA bacteriophage. Regarding the newly proposed framework to describe capsid organization, it appears to have T_t_(2,1) = 7 organization[30] and has a minor coat protein that sits in hexavalent positions (Figure 4). Patience also has a decoration protein that links the minor coat proteins together and makes no contact with the major capsid protein (Figure 5). The major capsid proteins that make up the hexamers (Figure 4) have an arrangement that appears similar to the 80alpha bacteriophage procapsid (EMD-7030) [31]. The internal diameter of the capsid is 63.5 nm.

**Figure 4.**
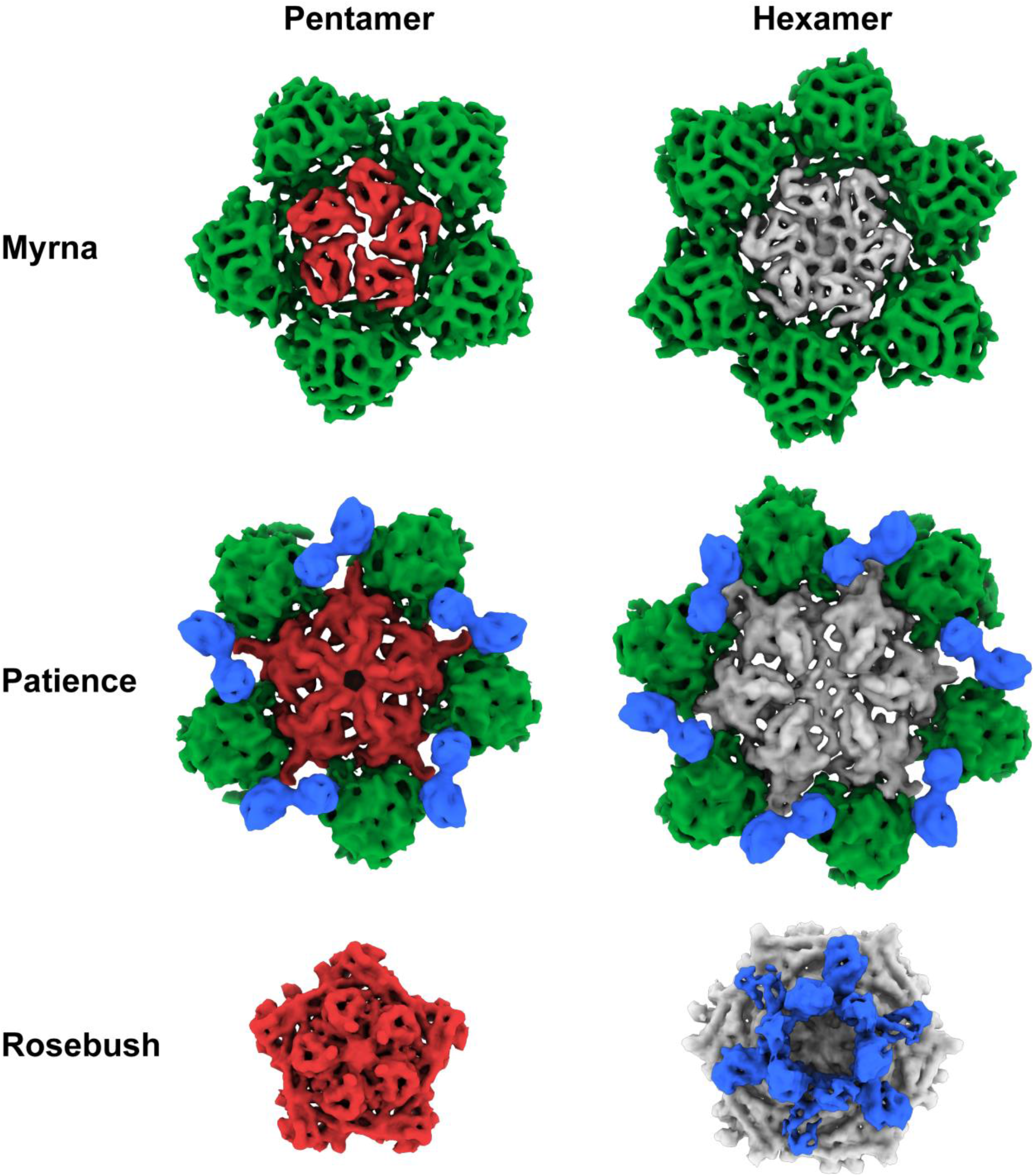
Hexamers and pentamers of the actinobacteriophages. The pentamers and hexamers (including minor/decoration proteins) have been segmented out from the capsid maps using the Segger[15] setting in Chimera[13]. Major capsid proteins are colored grey (hexamer) or red (pentamer). Minor capsid proteins green and decoration proteins blue. Images have not been re-scaled and are comparable.

**Figure 5.**
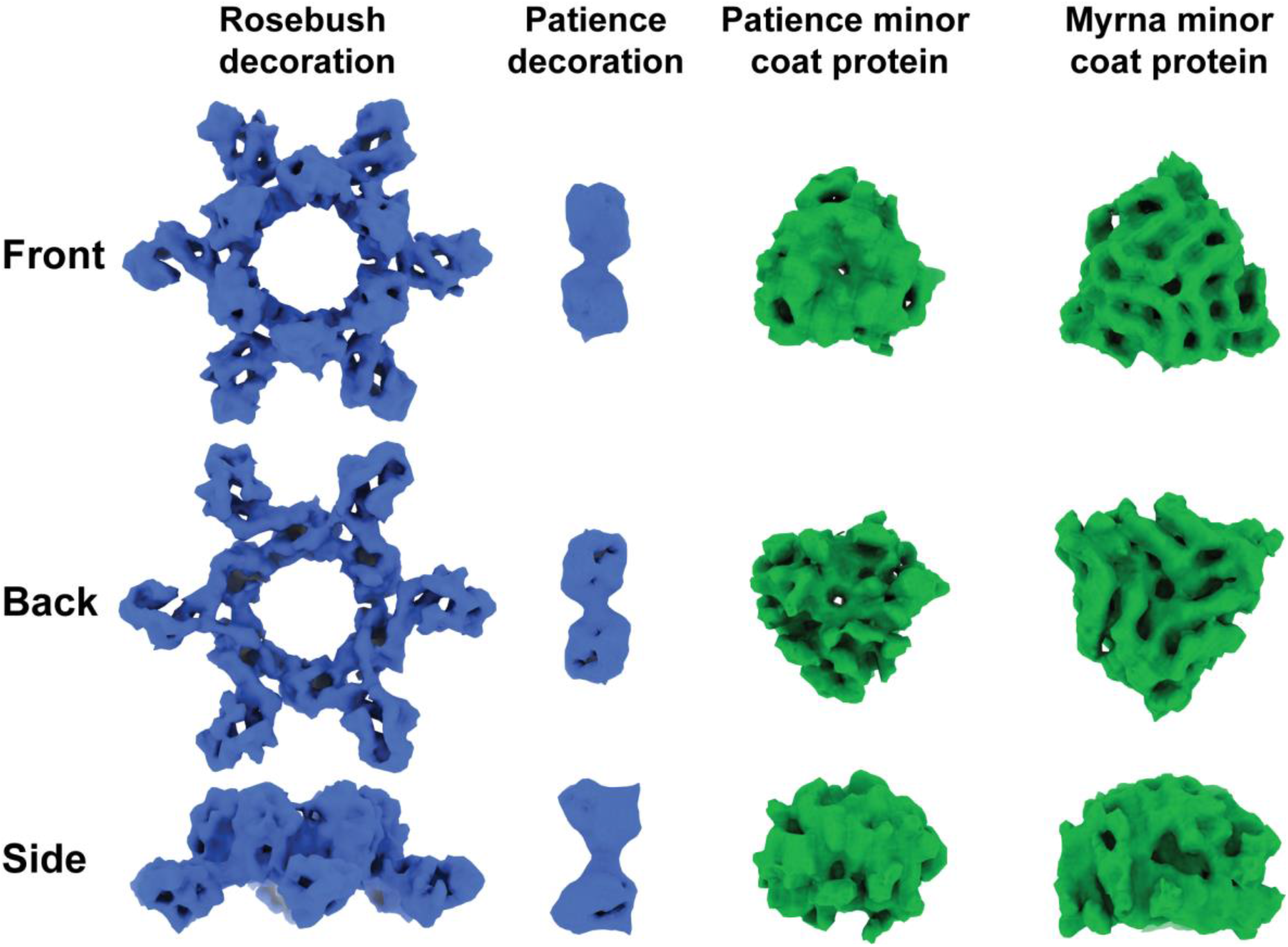
Density of segmented auxiliary proteins. Decoration proteins and minor protein density segmented from capsid maps using the Segger[15] setting in Chimera[13]. Front view is from outside of the capsid looking in. Back view is from the inside of the capsid looking out. Densities have been scaled relative to one another and their sizes are comparable.

Previous work characterized the proteins in the mature capsid of Patience using mass spectrometry[21] and SDS-PAGE. Their SDS-PAGE analysis showed three prominent bands that matched the predicted molecular weights of the capsid subunit (gp23), major tail subunit (gp31) and a protein with unknown function (gp15). The mass spectrometry analysis of the whole capsid showed that the top five most prevalent proteins in the capsid were the capsid subunit (gp23), gp15, gp4 and gp29 (gp31 was still identified but is not in the top five most prevalent).

Based on these previous results, we considered gp4, gp15 and gp29 (Figure 6) as candidates for the minor capsid and decoration protein based on the number of expected copies of each protein in the mature capsid. All three proteins had their structure predicted using the I-TASSER server[16,17] to enable fitting into the cryoEM density. Gp4’s position outside of the expected gene order (synteny) of structural proteins and small size (102 amino acids, 10.5 kDa) in the genome makes it unlikely to be part of the structural proteins. Gp29’s (139 amino acids, 15 kDa) predicted structure fits the density of the decoration protein well. However, the ITASSER C-score (−3.78) and TM-score (0.3 ± 0.1) suggest that the prediction is not that accurate. However, it’s position in the genome, and proximity to the major capsid protein (gp23) makes this protein most likely to be the decoration protein. The cryo-EM map, when fit with the ITASSER model (not shown), suggests that gp29 exits as a dimer in the capsid. With respect to the minor capsid protein, gp15 is the only remaining candidate based on the previous SDS-PAGE and mass spectrometry analysis. However, the I-TASSER model does not resemble the density, nor does HHPRED[18] and PSI-BLAST[19] predict any structural homology for any other known proteins.

**Figure 6.**
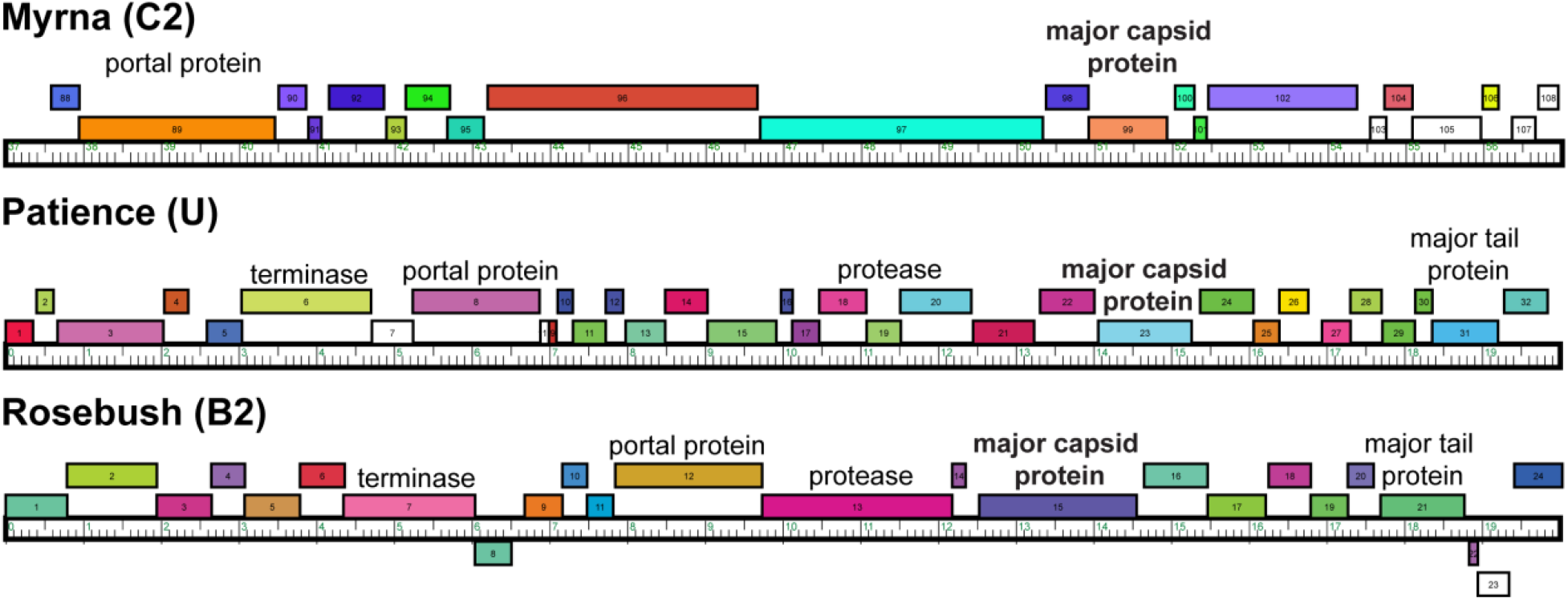
Gene organization of structure proteins. A portion of each genome displayed with the phamerator software[43] showing the positions of the annotated structural proteins.

**Rosebush** shows T=9 capsid organization. It has a decoration protein that forms raised hexamers that sit directly above the capsid hexamers and makes contacts with the major capsid proteins beneath. It has a similar internal capsid diameter to Patience (61.4 nm). The major capsid protein shows HK97-fold like properties, with the A domain and E-loop visible. However, HHPRED and PSI-BLAST fail to predict the HK97-fold, as does the I-TASSER software, suggesting that this is a novel variant of the HK97-fold. The major capsid protein is also relatively large at 71 kDa, with no obvious scaffolding protein upstream of gene *15* (Figure 6). The large size of the major capsid protein suggests it has a similar assembly pathway as HK97 with a combined scaffold/major capsid protein that is then cleaved after assembly.

**Myrna** shows T=16 capsid organization, with a minor capsid protein that sits in hexavalent positions. Following the new framework it is a T_t_ (4,0) = 64/3, similar to HSV-1[32]. The interior capsid diameter is 81.1 nm. The minor capsid protein gene *(98)* could be identified easily using bioinformatic analysis and sits directly upstream (Figure 6) of the major capsid protein gene (99), following the typical gene synteny found in *Caudovirales*. HHPRED suggests protein gp98 has structural homology (85.21% probability) with the TW1[33] bacteriophage (which infects *Pseudoalteromonas phenolica*, a gram-negative marine bacterium[34]) despite only having 20% amino acid sequence identify.

A related gene (based on amino acid sequence identity) can be found in the AA (host: *M. smegmatis)*, C1 *(M. smegmatis)* and DO *(G. terrae)* clusters, as well as the singleton Finch *(R. erthropolis)*, all of which have *Myoviridae* morphology. The other *Myoviridae* clusters DQ *(G. terrae* 3612), EA5 *(M. foliorum)*, AR *(Arthrobacter)*, AO1 *(Arthrobacter)*, and AO2 *(Arthrobacter)* do not have any gene with significant amino acid sequence identity to gp98. However, decoration proteins that have been annotated in these phages, for example protein 11 in Chipper1996 is predicted by HHPRED to also have structural similarity to the TW1 minor coat protein, suggesting a conserved fold in these minor coat proteins.

### SDS-PAGE gel and mass spectrometry analysis of Rosebush

Purified Rosebush was run on an SDS-PAGE gel and the four darkest bands were excised for mass spectrometry analysis (Figure 7 and Tables S1, S2 and S3). The rationale being that the number of each accessory protein in the capsid should be present in relatively high amounts and produce a dark band on the gel. One of the bands from the Rosebush gel (band 4 in Figure 7) was identified by mass spectrometry analysis to be gp17 (Figure 6). We propose that gp17 is the decoration protein.

**Figure 7.**
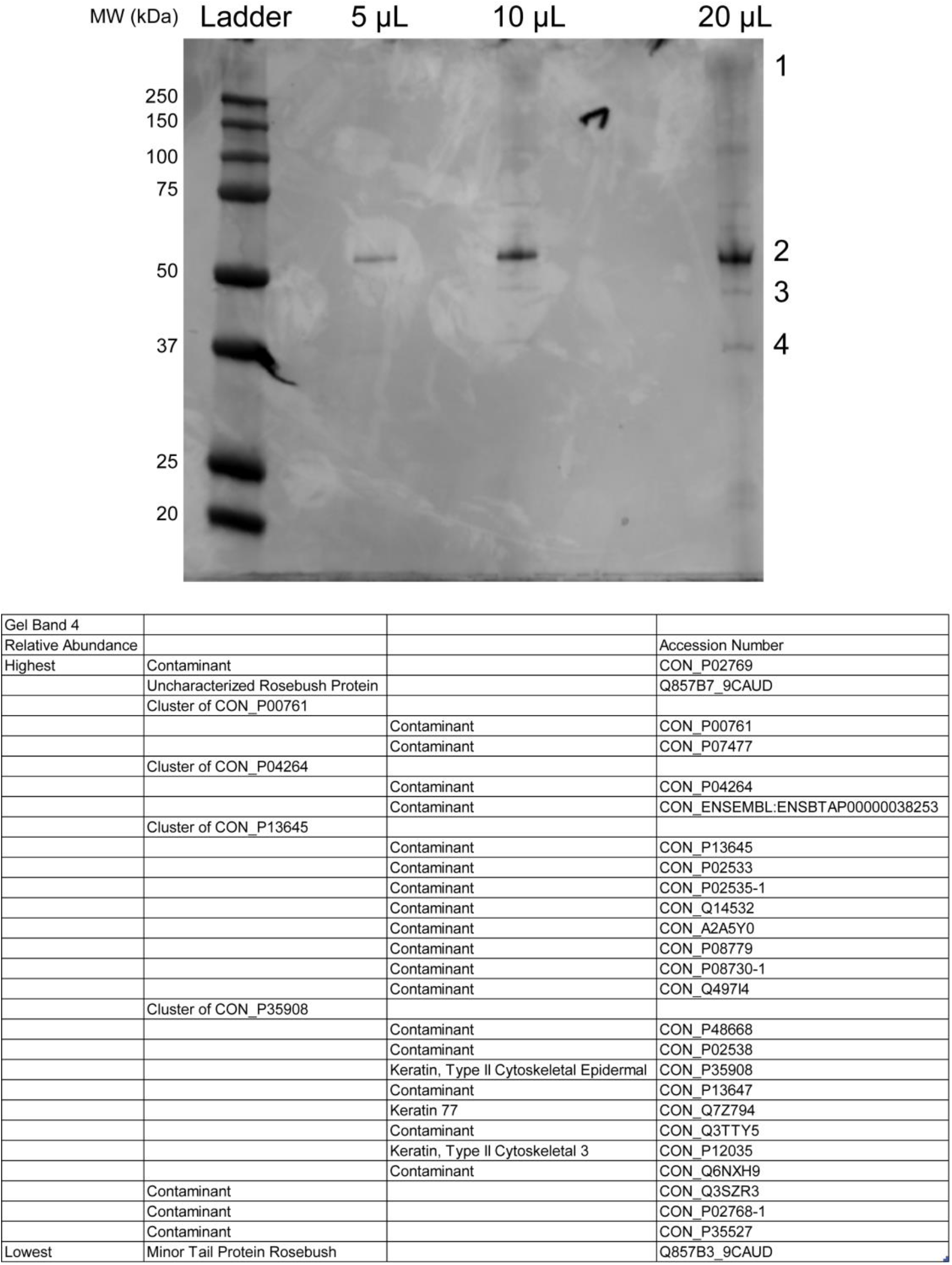
Mass spectrometry identification of accessory proteins in Rosebush. SDS-PAGE gel of Rosebush (top) with the bands excised for mass spectrometry highlighted by 1-4. Bottom: the results of the mass spectrometry shown for band 4 highlighting the phage proteins and contaminants identified.

### Bioinformatic analysis of accessory protein gp17

BLAST-P[19] of the amino acid sequence of gp17 against the non-redundant protein sequences database shows that gp17 is found in a number of other actinobacteriophages in Cluster B. Those phages in the B2 sub-cluster have almost 100% sequence identity to Rosebush’s gp17 while those phages in other B sub-clusters have far lower sequence identity; for example phage Phelemich[4] in the B5 sub-cluster has a 31.58% amino acid sequence identity. Partial matches can also be found in the DR cluster (6 members); for example gp15 of CloverMinnie, which infects the *G. terrae* host. TPA2 (a singleton that infects *Tsukamurella paurometabola* Tpau37) also shows a partial match. HHPRED and PSI-BLAST analysis shows no significant matches to any known protein structure. The amino acid sequence was submitted to the I-TASSER protein structure prediction software, but the resulting models (not shown) had C-scores and TM-scores which suggested a poor predicted model (the best model had a C-score of −3.98 and a TM-score of 0.29 ± 0.09). Comparison of the models to the cryo-EM map showed a poor fit.

## Discussion

The phage structures described here expand our structural knowledge of bacteriophages and their diversity. The decoration proteins of Patience (gp29) and Rosebush (gp17) are novel and have no known structural homologues. Their purpose is unknown and further experiments are needed, for example deletion of the decoration proteins from the phages, to measure their effect on capsid stability and infectivity. We hypothesize that both accessory proteins of Patience (which we annotate as gp15 and gp29) are likely involved in capsid stability with gp29 acting like other minor coat proteins and gp15 linking them together. The role of gp17 in Rosebush is likely to have a minimal effect on structural stability based on the observation that it extends far from the phage capsid and makes few contacts with the coat protein hexamer underneath. It may play a role in host recognition, making weak contacts with the host before the tail contacts its receptor.

Patience and Rosebush are excellent examples of two of the main mechanisms by which bacteriophages can increase the size of their capsid while using the same HK97-fold building block. Both Patience and Rosebush have similar capsid sizes (63.5 and 61.4 nm internal diameter, respectively) but different T numbers. Rosebush has altered its major capsid protein, and presumably the scaffolding protein that these bacteriophages use in the capsid assembly process. The altered major capsid protein is used to increase the T number from 7 to 9. This change in T number causes the number of major capsid proteins to increase from 420 copies to 540, resulting in a larger capsid. Patience has used a completely different method to increase capsid size by using the same number of major capsid proteins (420 copies in a T=7 capsid), but adding minor capsid proteins to increase the capsid size. The minor capsid protein of Patience may also play a role in stability in a similar way to the minor capsid protein gpD in phage lambda[35]. In phage lambda, the position of gpD in the capsid is where the cross-link can be found in the HK97 bacteriophage major capsid protein, which is important in its capsid stability. It is thought that gpD minor capsid protein replaces the need for the major capsid protein cross-linking.

Structural characterization of T=9 viral capsids is relatively rare with only Basilisk (Bacillus cereus, 18 Å resolution)[35], Ty3/Gypsys retrotransposon capsid (7.5 Å resolution, 6R24)[36], *H. ochraceum* microcompartment shell (3 Å resolution, 6MZX)[37,38], and N4 (14 Å resolution)[39] having published structures, all at a lower resolution than Rosebush described here. Rosebush’s coat protein configuration appears completely different to other bacteriophages. Further structural prediction and comparison to known databases reveals that the Rosebush major capsid protein is a novel variant of the HK97-fold. Further work is needed to obtain a higher-resolution map to allow for amino acid model building.

The structure of Myrna shows how common the T=16 capsid architecture is, with examples of viruses found in all domains of life that have a similar structure, for example the human virus HSV-1[32]. This is the first time it has been described in the Actinobacteria, and suggests that this capsid architecture is adaptable to a number of different hosts and environments to encapsulate large genomes.

Finally, we have shown that the use of multiplexing phage capsids is a successful strategy for lowering the cost per structure. Mixing different phage capsids together allows a single data collection session on expensive cryo-electron microscopes. We predict that up to five phages could be easily multiplexed on a single grid, further decreasing the cost. This is an important tool for viral structural studies. There are over 100 sub-clusters in the actinobacteriophage alone. In order to quickly, and relatively cheaply, screen a phage from each sub-cluster, we believe that multiplexing is the best method. Once all the phages have been screened at 6 angstrom resolution, which allows for the HK97-fold to be identified, they can be classified into structural groups and exemplar phage identified for high resolution structural studies. This method could be widely used for phages that infect various phyla to rapidly expand the structural capsid database to allow comparison of phage. This will have the greatest impact on viral evolution studies. It has long been discussed in the literature that the major capsid protein may be the best way to explore the evolution of viruses[40–42]. However, the conservation of the protein may only be found in the protein fold and not in the amino acid sequence. Therefore, collecting many phage structures is the only way to carry out this evolutionary work.

## Conclusions

We have identified four head accessory proteins in the actinobacteriophages, two are novel decoration proteins and two are minor capsid proteins that show structural homology to known minor capsid proteins in other viruses.

## Acknowledgements

We would like to thank the University of Massachusetts cryo-EM core facility and their directors Dr Chen Xu, and Dr KangKang Song as well as their staff Dr Kyounghwan Lee for their help in cryo-EM data acquisition.

We gratefully acknowledge the quantitative proteomics analysis conducted by Dr. Jeremy L. Balsbaugh, Director of the Proteomics & Metabolomics Facility, and Zachary Cosgrove. The facility is a component of the Center for Open Research Resources and Equipment at the University of Connecticut.

## Data deposition

EMDB accession numbers are as follows:

Rosebush: EMD-21122, Patience: EMD-21123, Myrna: EMD-21124

