## Supplemental Tables and Figures for "Structures of three actinobacteriophage capsids: Roles of symmetry and accessory proteins"

**Supporting information**

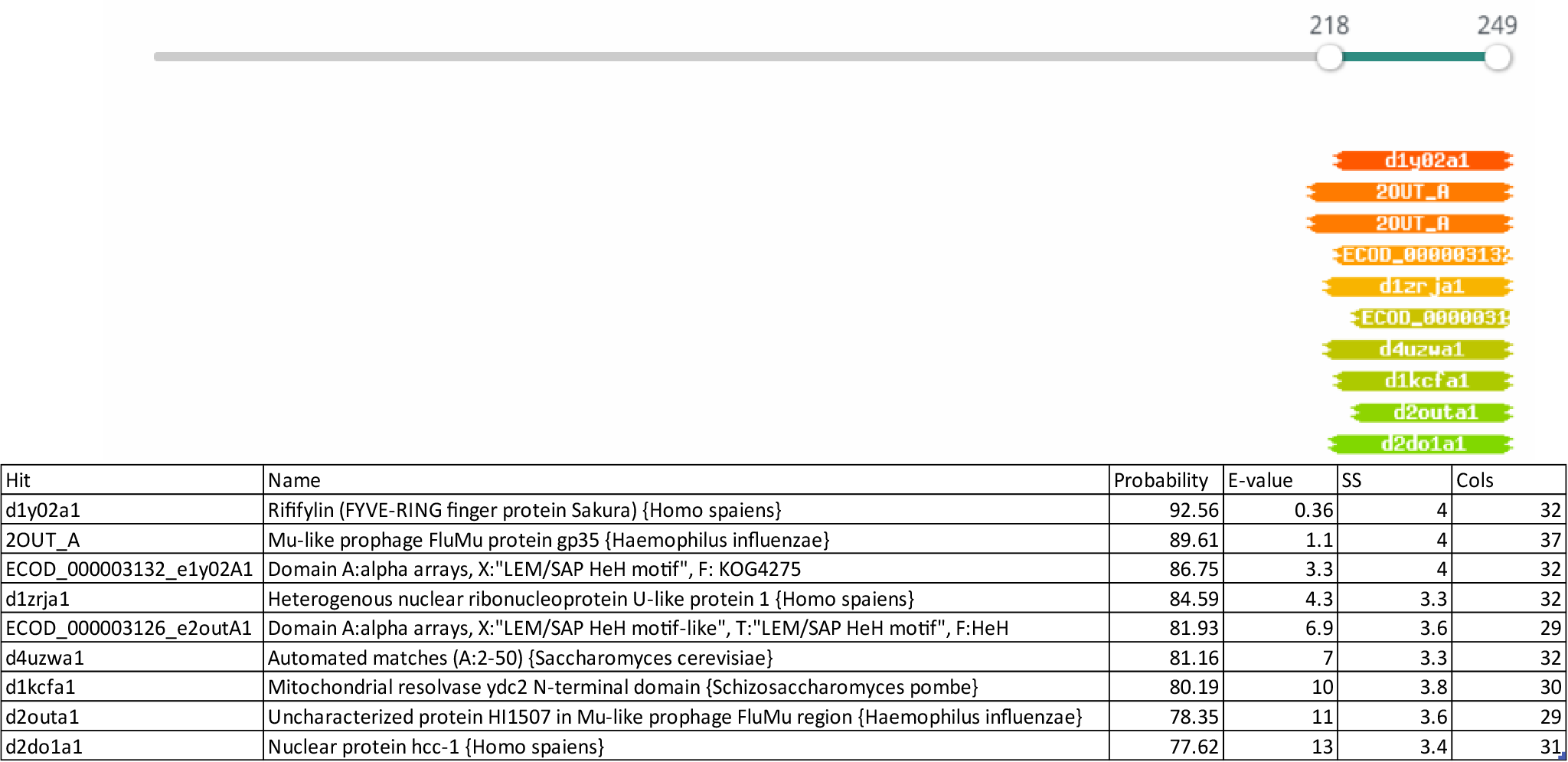

**Supporting Figure S1. HHPRED prediction of Rosebush GP17.** The amino acid sequence of GP17 was used as the input into HHPRED and searched against all the available databases.

**Table S1. Mass spectrometry data analysis of band 1 from SDS-PAGE of Rosebush capsid**

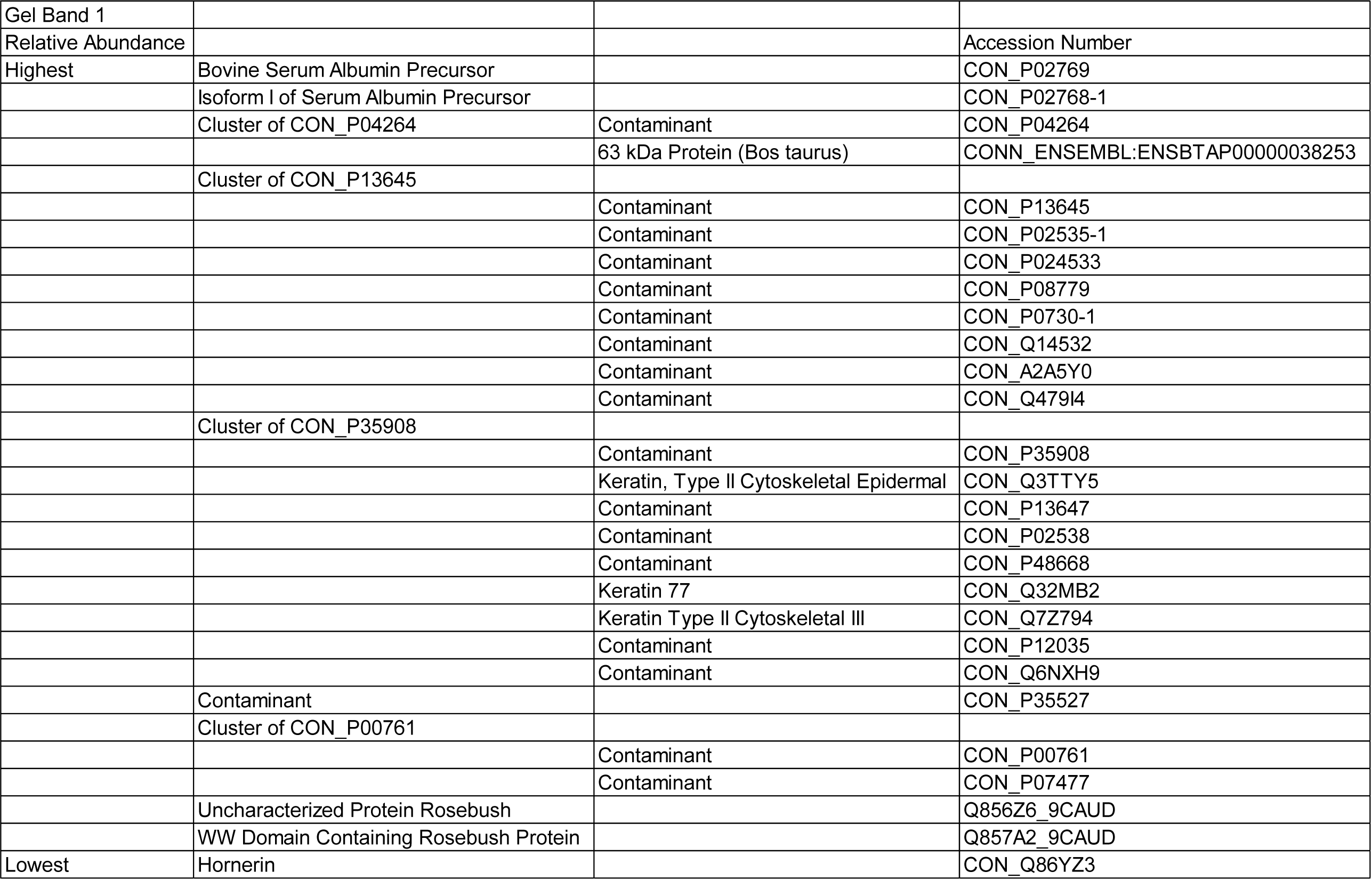

**Table S2. Mass spectrometry data analysis of band 2 from SDS-PAGE of Rosebush capsid**

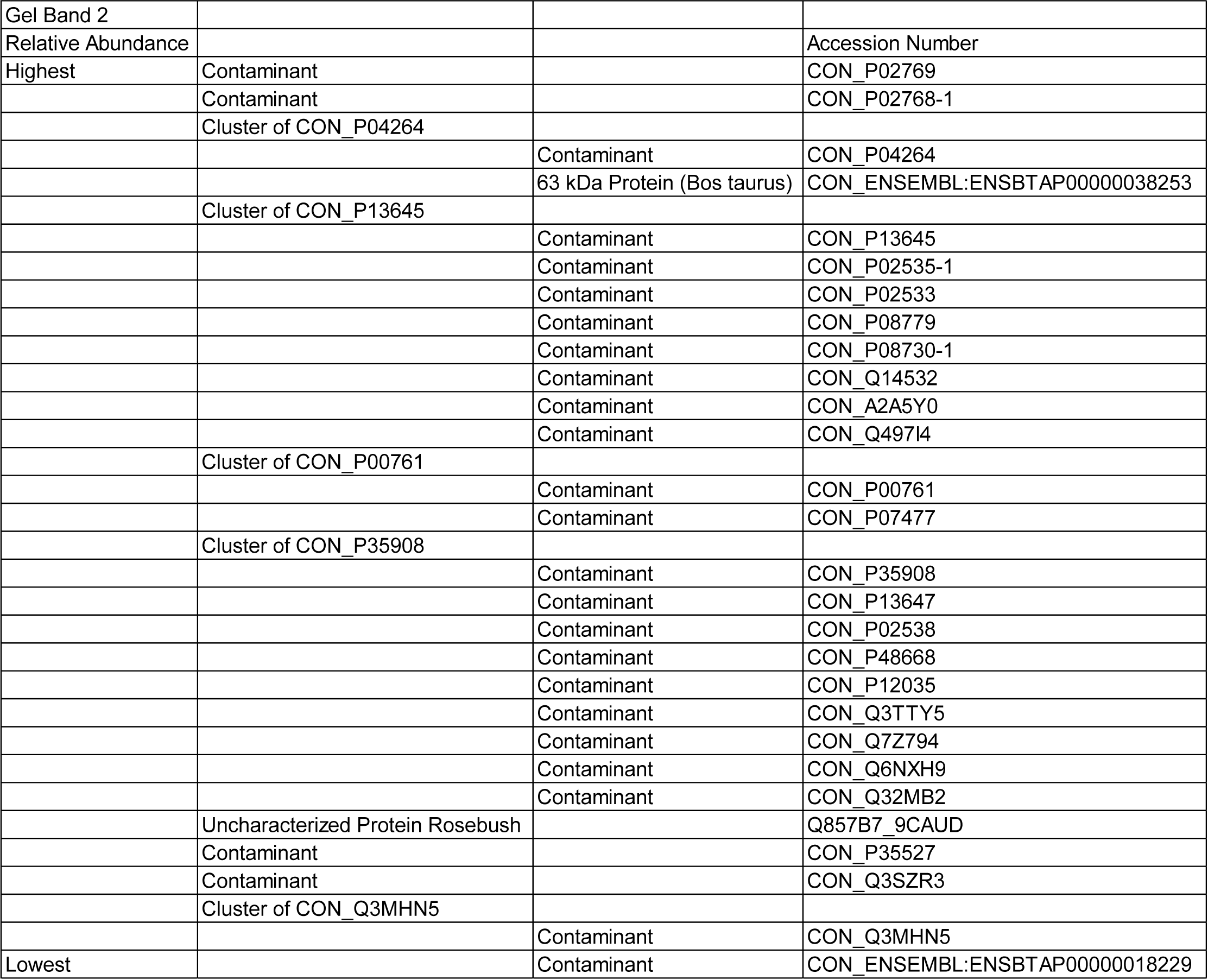

**Table S3. Mass spectrometry data analysis of band 3 from SDS-PAGE of Rosebush capsid**

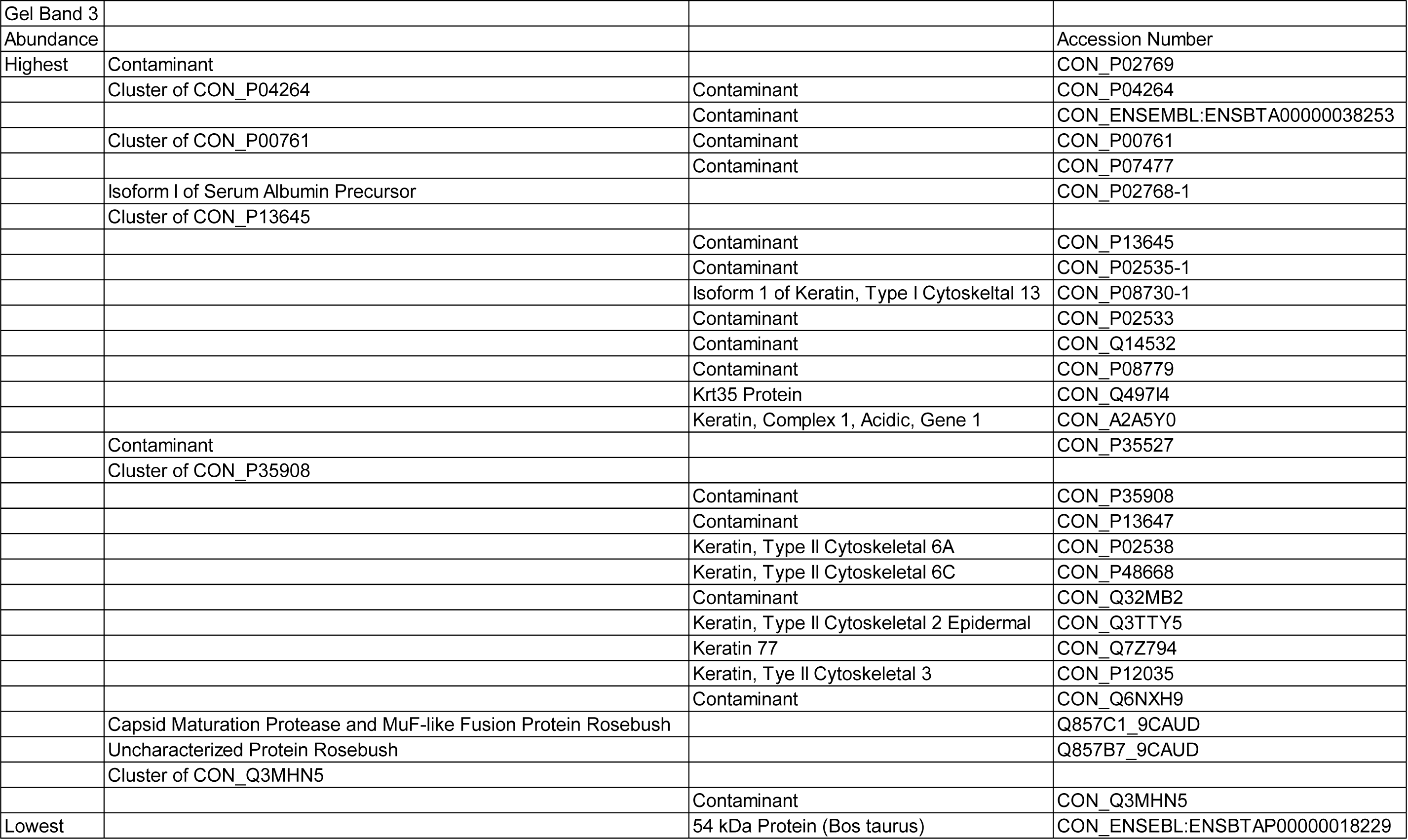
